# Detection of the 40-Hz Auditory Steady-state Response with Optically Pumped Magnetometers

**DOI:** 10.1101/2021.10.01.462598

**Authors:** Kyung-min An, Jung Hyun Shim, Hyukchan Kwon, Young-Ho Lee, Kwon-Kyu Yu, Moonyoung Kwon, Woo Young Chun, Tetsu Hirosawa, Chiaki Hasegawa, Sumie Iwasaki, Mitsuru Kikuchi, Kiwoong Kim

## Abstract

Magnetoencephalography (MEG) is a functional neuroimaging technique that noninvasively detects the brain magnetic field from neuronal activations. Conventional MEG measures brain signals using superconducting quantum interference devices (SQUIDs). SQUID-MEG requires a cryogenic environment involving a bulky non-magnetic dewar and the consumption of liquid helium, which restricts the variability of the sensor array and the gap between the cortical sources and sensors. Recently, miniature optically pumped magnetometers (OPMs) have been developed and commercialized. OPMs do not require cryogenic cooling and can be placed within millimeters from the scalp. In the present study, we arranged six OPM sensors on the temporal area to detect auditory-related brain responses in a two-layer magnetically shielded room. We presented the auditory stimuli of 1-kHz pure-tone bursts with 200-ms duration and obtained the M50 and M100 components of auditory evoked fields. We delivered the periodic stimuli with a 40-Hz repetition rate and observed the gamma-band power changes and inter-trial phase coherence of auditory steady-state responses at 40 Hz. We found that the OPM sensors have a performance comparable to that of conventional SQUID-MEG sensors, and our results suggest the feasibility of using OPM sensors for functional neuroimaging and brain–computer interface applications.

## 1. Introduction

An auditory steady-state response (ASSR) is the result of the entrained neural rhythm in the primary auditory region generated by the periodic repetition of an auditory stimulus (Hari et al., 1989). In humans, the ASSR is known to have a maximum magnitude at approximately 40 Hz, which is the resonance frequency of the auditory neural circuit (Galambos et al., 1981; Pastor et al., 2002; Ross et al., 2000). The amplitude and phase of 40-Hz ASSR are supposed to reflect the balance between the inhibitory GABAergic and excitatory glutamatergic neurons (Sivarao et al., 2016; Tada et al., 2020).

Two methods have been used to investigate 40-Hz ASSR, namely the event-related spectral perturbation (ERSP) and inter-trial phase coherence (ITPC) methods. The ERSP is a measure of induced power changes and is independent of the phase. The ITPC is a measure of the phase synchronization across trials and is also called the phase-locking factor (Delorme and Makeig, 2004; Tallon-Baudry et al., 1996).

Reduced power and phase synchronization of 40-Hz ASSR have been reported in individuals with schizophrenia (Tada et al., 2020; Thune et al., 2016), bipolar disorders (O’Donnell et al., 2013; Rass et al., 2010), and autism spectrum disorders (Ono et al., 2020; Seymour et al., 2020).

The 40-Hz ASSR can be non-invasively measured using scalp electroencephalography (EEG) and magnetoencephalography (MEG). EEG and MEG measure neurophysiological activities with high temporal resolution. EEG has the advantages of a relatively simple and cost-effective system and the flexible arrangement of sensors. However, EEG has a longer preparation time for attaching electrodes on the scalp and has limited spatial resolution owing to the low and inhomogeneous electrical conductivity of the skull (Srinivasan et al., 1998). MEG has a high spatial resolution because the neuro-magnetic field is not sensitively affected as it passes through head tissue (Hamalainen et al., 1993; Stenroos and Nummenmaa, 2016). Conventional MEG measures magnetic fields generated by neurons using a superconducting quantum interference device (SQUID). Low-temperature SQUID sensors usually operate at approximately 7 K with the use of liquid helium. A rigid reservoir for the liquid helium is required to maintain a cryogenic temperature. The use of a rigid reservoir requires the SQUID sensors to be fixed inside a helmet, and the distance between the sensors and scalp is at least approximately 2 cm.

Recently, optically pumped magnetometers (OPMs) with a small size of 12.4 mm × 16.6 mm × 24.4 mm have been developed and commercialized (Budker and Romalis, 2007; Shah and Wakai, 2013). The OPM sensor operates at room temperature and can be placed close to the scalp in a flexible manner. OPM sensors have been applied to detect neuromagnetic signals with such advantages. Previous OPM-based MEG studies have measured various brain activities relating to auditory evoked fields (AEFs) (Borna et al., 2020; Borna et al., 2017; Kim et al., 2014; Kowalczyk et al., 2021), visual processing (Iivanainen et al., 2020), somatosensory processing (Boto et al., 2017), motor processing (Boto et al., 2018), and language function (de Lange et al., 2021; Tierney et al., 2018).

There is, however, no report of the OPM-MEG measurement of 40-Hz ASSR, which reflects the functions of gamma-band activity and has potential clinical application. In this study, we developed an OPM-MEG system using six OPM sensors to detect auditory brain responses from the temporal lobe. We presented participants with auditory pure-tone bursts while conducting OPM-MEG recordings and confirmed that the OPM sensors can detect the AEFs. Additionally, we delivered repetitive auditory stimuli at 40 Hz and demonstrated that the OPM can reliably detect 40-Hz ASSR by calculating the ERSP and ITPC.

## 2. Materials and Methods

### 2.1. Participants

Twenty-two right-handed healthy participants (mean ± SD age, 27.05 ± 4.36 years; 11 females) participated in the study. Handedness was assessed using a translated version of the Edinburgh Handedness Inventory (Oldfield, 1971). All participants had normal hearing and normal or corrected-to-normal vision and none reported any neurological or psychiatric disorder. The experimental procedures were approved by the Ethics Committee of the Korea Research Institute of Standards and Science (KRISS-IRB-2021-04). All participants gave their written informed consent.

### 2.2. Experimental Paradigm and Stimuli

We presented two types of auditory stimulus during the OPM-MEG recordings. We first tested pure-tone auditory stimuli to confirm that OPM-MEG can detect the AEFs. The pure-tone stimulus was a 1-kHz tone burst with a duration of 100 ms. We delivered pure-tone bursts 230 to 240 times with an inter-stimulus interval of 1.8 to 2.3 s in one session.

We used auditory click-train sounds to elicit the ASSR at gamma frequency. The auditory click-train was created with 1-ms pulse sounds delivered at 40 Hz for 1 s. We presented a total of 250 click-train stimuli with an inter-stimulus interval of 2.5 to 3 s in two sessions. Each session lasted approximately 6 min.

During the recordings, participants stared at a fixation point and the auditory stimuli were presented at 80 dB to the right ear through a MEG-compatible ear tube. To help the participants concentrate on the task, the stimuli were randomly delivered, and we asked the participants to count the number of stimuli delivered in each session.

### 2.3. OPM-MEG acquisition

OPM-MEG was measured using an array of six OPMs (Gen-2.0 QZFM; QuSpin Inc., Louisville, CO). The measurement setup is shown in **Figure 1**. The OPM sensors were mounted on a three-dimensionally printed curved plate that fitted the temporal head surfaces of the participants. The plate had an arched hollow on the bottom side in which to place a participant’s left ear. This hollow made it comfortable for the participant to lean their head close to the sensor plate. The OPM sensor plate had nine sockets with separations of 15 mm in three row and three columns. The six OPM sensors were fixed in the sockets of the two lower rows (indicated by blue rectangles in Figure 1a). The center-to-center distance between adjacent OPM sensors was 31.6 mm horizontally and 27.4 mm vertically.

**Figure 1.**
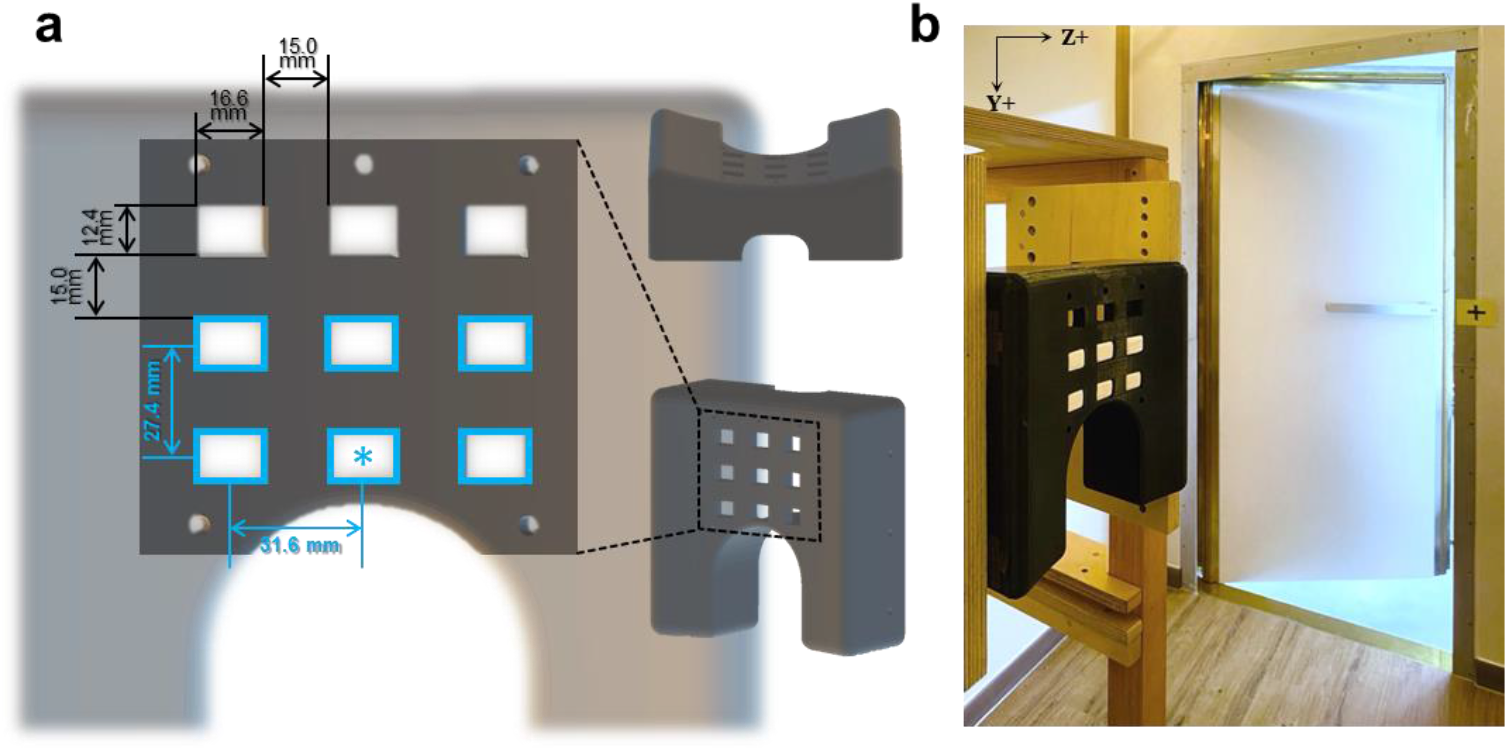
Optically pumped magnetometer (OPM) sensor array and measurement setup in the magnetically shielded room. **a**. We developed a sensor array for arranging six OPM sensors. The OPM sensor array had nine sockets in which to insert OPM sensors. Six OPM sensors were fixed in the sockets marked by blue rectangles. The center sensor (marked with an asterisk) was positioned on the participant’s T3 point, which was overlying the left auditory cortex according to the EEG sensor layout. **b**. The OPM sensor array was placed in the middle of the two-layer magnetically shielded room. The OPM sensor array was positioned to cover the temporal region of the participant’s left head while the participant was sitting in the magnetically shielded room.

To find the appropriate sensor position for detecting the auditory brain signals, we marked the T3 and Cz points of each participant according to the EEG 10-20 lead system. The T3 point is the scalp site overlying the left-hemisphere auditory area of the cerebral cortex. The Cz point is the midline central point of the scalp. We placed the center sensor (indicated by the blue asterisk in Figure 1a) on the T3 point of each participant and aligned the vertical axis of the sensor array along the line between T3 and Cz. The OPM sensor array were placed in the middle of the two-layer magnetically shielded room (Korea Research Institute of Standards and Science, Republic of Korea) (Figure 1b). During the OPM-MEG recordings, the participants were seated and leaned toward the sensor array and headrest such that their head was close to the sensors.

The electronics controller of the OPM system delivered two analogue outputs for the magnetic field strength in the y- and z-directions for each sensor. The analog signals and auditory trigger were simultaneously sampled by a 16-bit data acquisition system (NI-9205, National Instruments Co., Austin, TX) at a sampling rate of 1 kHz. The scaling of the output voltage to the measured magnetic field was 2.8 V/nT. We used only the signals of the z-direction in our analysis.

### 2.4. Data analysis

We analyzed the OPM-MEG data using the Brainstorm toolbox (Tadel et al., 2011), FieldTrip toolbox (Oostenveld et al., 2011), and MATLAB (The MathWorks). Raw data were bandpass filtered from 0.2 to 100 Hz. We applied a powerline notch filter at 60 Hz and band-stop filter at 21.5 and 27 Hz (±0.5 Hz), which are the frequencies of environmental vibration noise. We were segmented data from − 3 to 3 s following the onset of each auditory stimulus. We rejected the trials containing obvious artifacts over 300 fT.

To calculate the event-related fields, we applied a low-pass filter to the data of pure-tone bursts at 40 Hz and to the data of 40-Hz ASSR at 60 Hz. The individual AEFs were DC normalized with the baseline from normalized with the baseline −200 to 0 ms according to the auditory stimulus onset. We averaged AEFs within each participant and across all participants to obtain the grand-average AEFs. To obtain the field distribution for the AEFs of pure-tone bursts, we calculated magnetic field maps of the baseline (−110 to −100 ms according to the auditory stimulus onset), M50 (40 to 50 ms), and M100 (80 to 90 ms) components.

To calculate the power changes and phase synchronization of ASSR, we applied time-frequency analysis at 1–60 Hz using a seven-cycle Morlet wavelet for each single trial. The time-frequency representations (TFRs) were calculated by converting to the percentage changes in power relative to the baseline (−1.1 to −0.1 s). TFRs were averaged for each participant and then grand-averaged for all participants. We assessed the significant time-frequency component related to the ASSR by comparing with the baseline period (−1.1 to −0.1 s) applying a parametric t-test (two-tailed). A correction of the false discovery rate was applied to control for type I error in the t-test. The alpha level was set at 0.05 in the statistical analysis. We calculated the gamma-band response modulated at 40 Hz by averaging the power changes from 38 to 42 Hz.

After decomposing the clean trial data using Morlet wavelets at 1 to 60 Hz, we calculated the ITPC as (Delorme and Makeig, 2004)

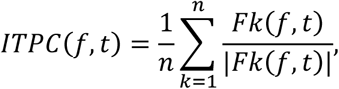

where *t* is time, *f* is the frequency, *n* is the number of trials, and *Fk*(*f, t*) is the spectral estimate of trial *k* at frequency *f* and time *t*. The ITPC reflects the phase synchronization at each time-frequency point. ITPC values range from 0 to 1 for a given frequency and time point. Larger ITPC values represent higher consistency in the phase synchronization and smaller ITPC values represent lower phase synchronization across trials (Cohen, 2014).

## 3. Results

We first delivered pure-tone bursts and calculated AEFs to confirm that our OPM sensor array could detect brain auditory activities. We observed clear AEFs detected by the six OPM sensors. Figure 2a presents the sensor distributions of the grand-average AEFs across the 22 participants. We obtained maximum activities of AEFs from the center sensor, which covered the T3 scalp site. Figure 2b shows the clear M50 and M100 components. The M50 component appeared at around 43 ms (42.91 ± 6.12 ms) whereas the M100 component appeared at around 86 ms (86.27 ± 7.34 ms). Figure 2c presents the topological map patterns of the baseline period (−110 to −100 ms according to the auditory stimulus onset), M50 component (40 to 50 ms), and M100 component (80 to 90 ms). The topologies of the M50 and M100 components had opposing polarity.

**Figure 2.**
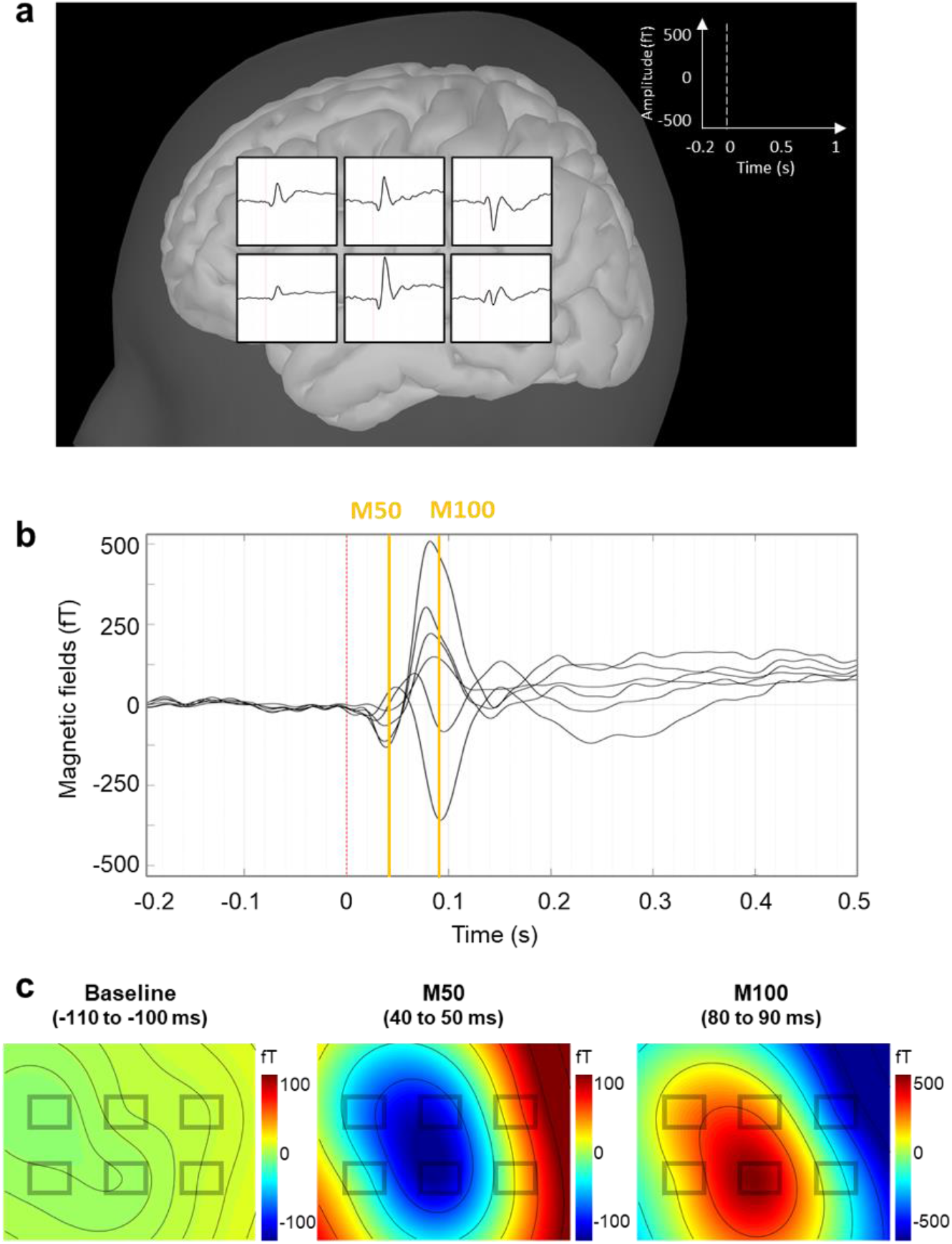
Auditory evoked fields and their topographical distributions recorded by OPM sensors during auditory pure-tone bursts. **a**. Grand-average auditory evoked field of 22 participants measured by each OPM sensor. **b**. M50 component observed at around 43 ms (42.91 ± 6.12 ms) and M100 component observed at around 86 ms (86.27 ± 7.34 ms). **c**. Topographical maps representing the field distributions of the baseline (−110 to −100 ms according to the onset of the auditory pure-tone burst), M50 component (40 to 50 ms), and M100 component (80 to 90 ms).

We presented auditory click-train stimuli at 40 Hz to the participants to investigate the modulated auditory gamma-band activity. The duration of the auditory click-train stimuli was 1 s. We calculated event-related fields, TFRs, and ITPC, to investigate the 40-Hz ASSR.

Figure 3a shows the grand-average AEFs related to the repetitive auditory stimuli at 40 Hz. We observed the M50 and M100 components at the early response and found brain waveforms modulated at 40 Hz lasting for 1 s.

**Figure 3.**
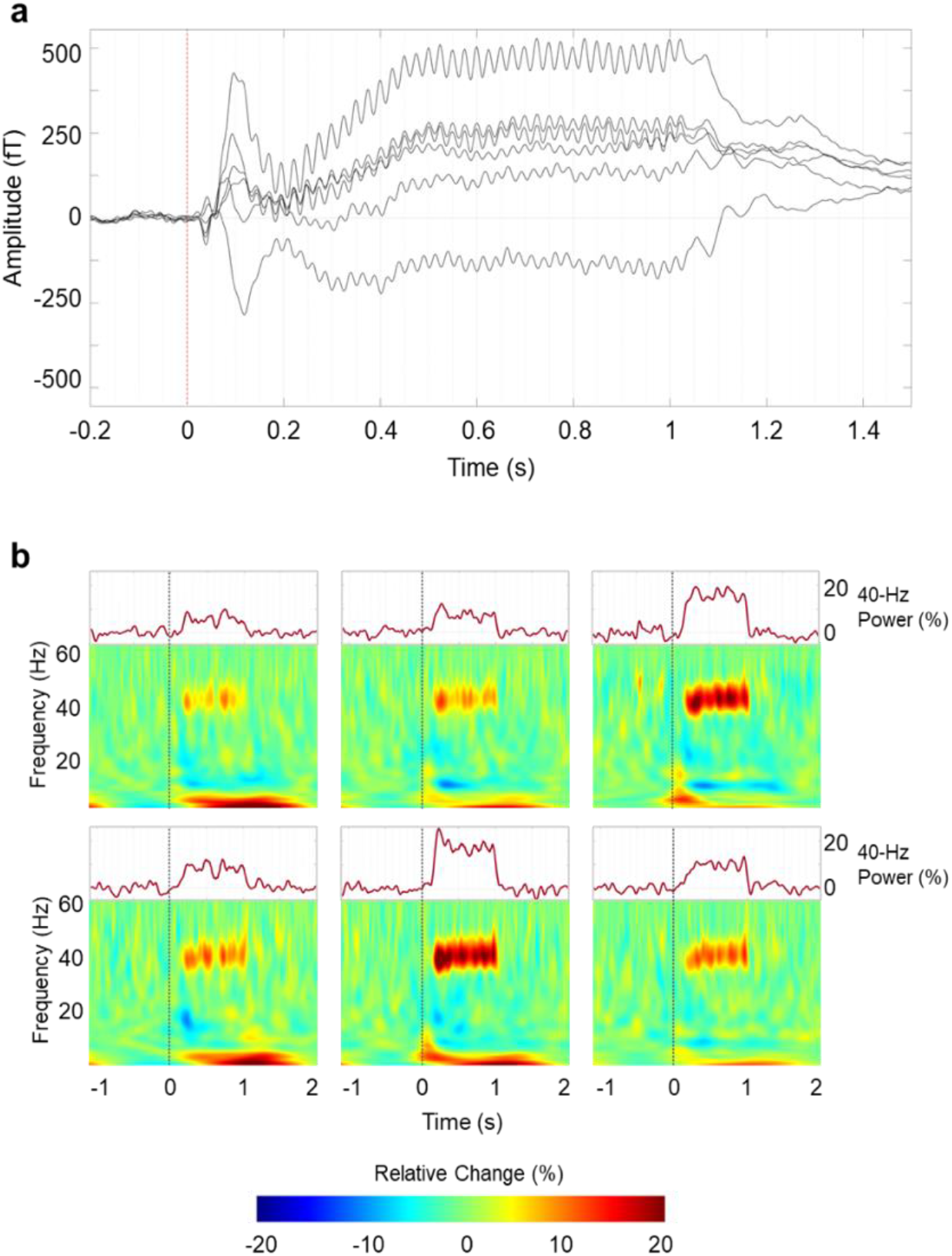
Grand-average waveforms and time-frequency representations during the 40-Hz auditory steady-state response. **a**. Grand-average waveforms recorded by the six OPM sensors show that the magnetoencephalographic fields were modulated by the repetitive auditory stimuli at 40 Hz. **b**. Grand-average time-frequency representations show that the relative power increased in the 40-Hz gamma band. Upper panels show the mean power changes of the gamma frequency band at 38 to 42 Hz. The 40-Hz gamma power increased for 1 s when 40-Hz auditory steady-state response stimuli were presented.

Group-averaged TFRs for the repetitive auditory stimuli at 40 Hz are plotted for each OPM sensor (lower panel of Figure 3b). The power of the gamma-band response increased at 40 Hz for about 1 s. The upper panel of Figure 3b shows the power changes of 40-Hz ASSR obtained by averaging from 38 to 42 Hz. We see that the gamma-band responses increased during the presentation of the 40-Hz auditory click-train stimuli (Figure 4).

**Figure 4.**
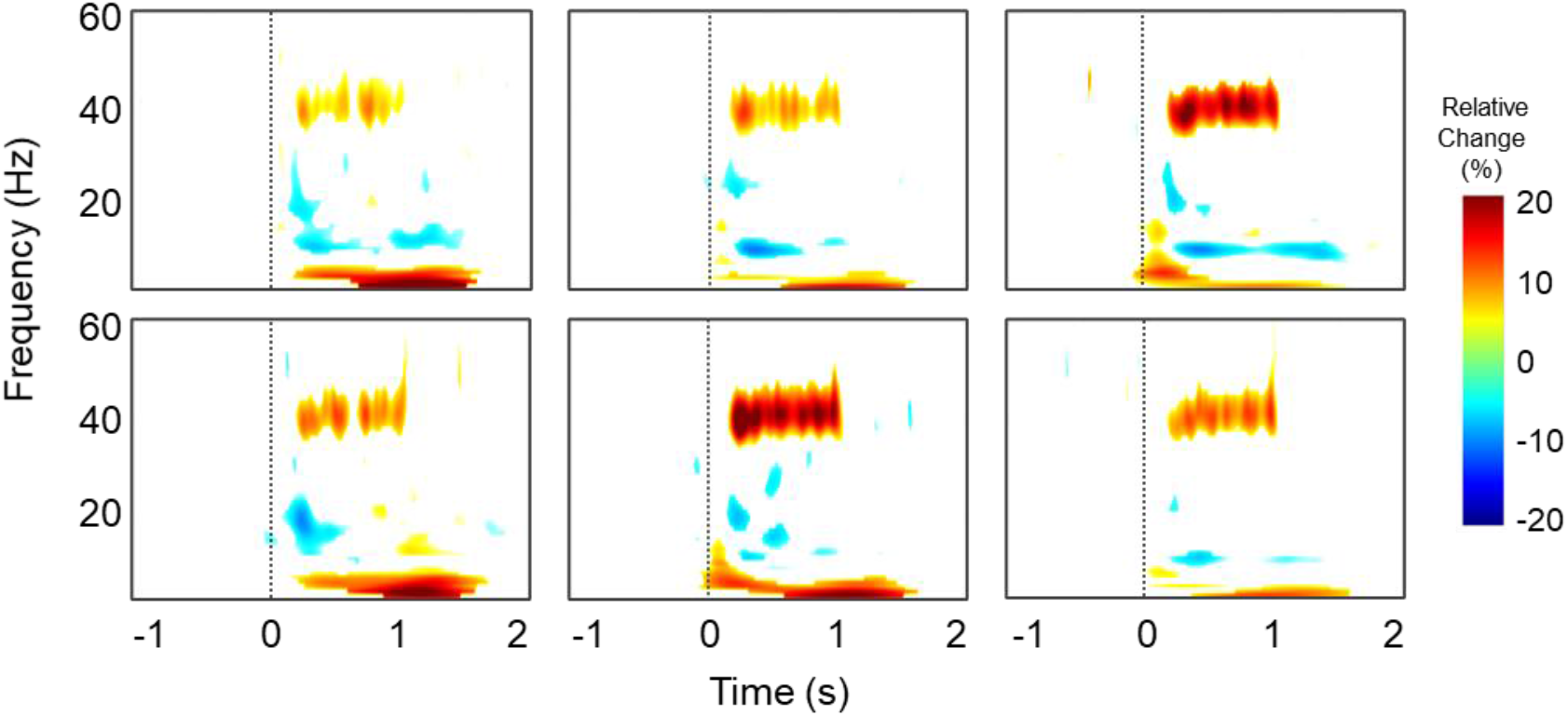
Grand-average time-frequency maps of statistical t-values for each OPM sensor. The maps show the power changes of statistically significant differences relative to the baseline period (−1.1 to −0.2 s) obtained in a parametric t-test with a correction of the false discovery rate for multiple comparison.

We analyzed the ITPC to investigate the phase synchronization of 40-Hz ASSR. Figure 5 shows the results of the ITPC for each OPM sensor. We find strong phase-locking at 40 Hz across trials. All our results show a maximum value for the center sensor, which overlaid the T3 point.

**Figure 5.**
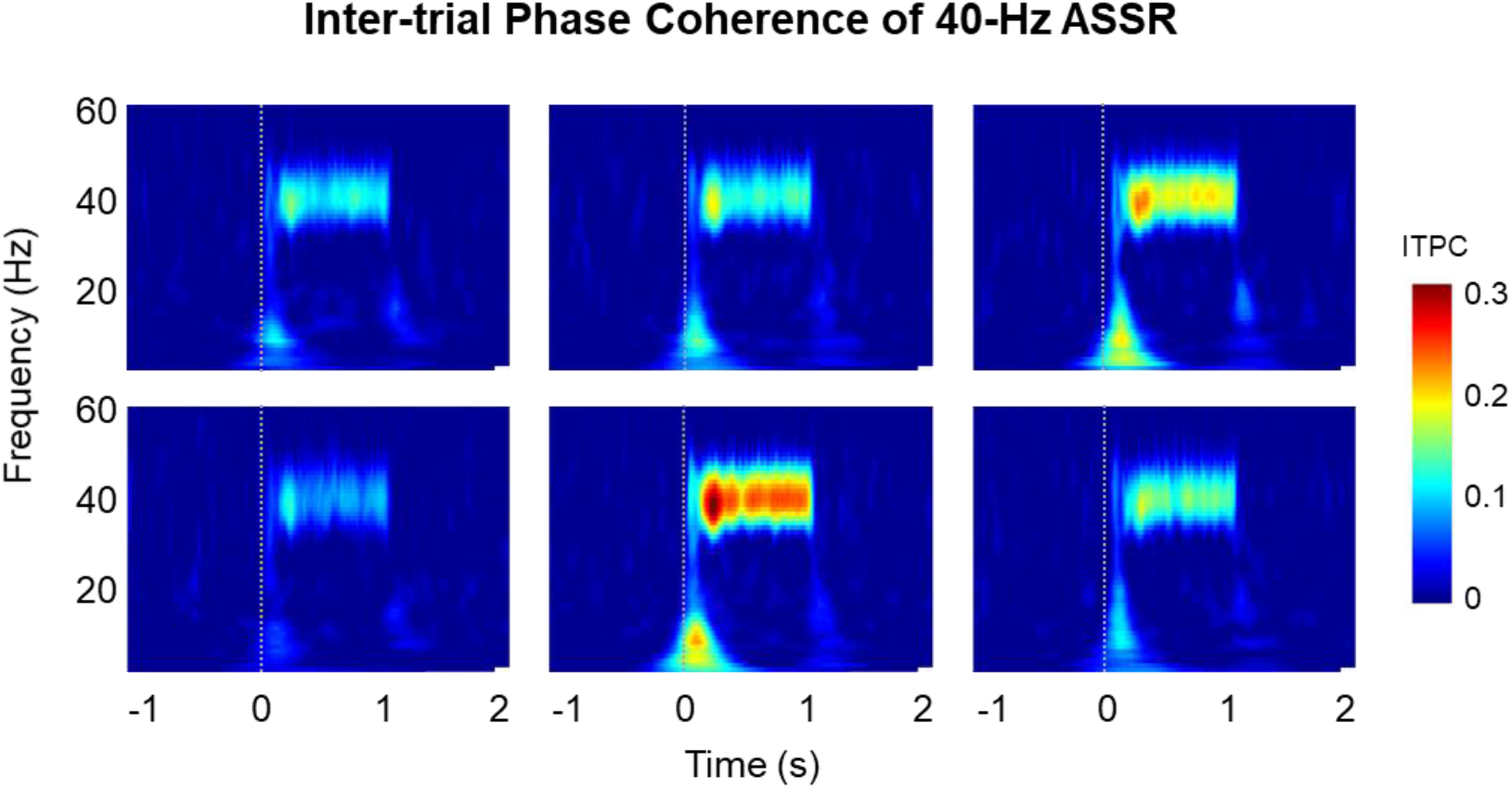
Results of inter-trial phase coherence of 40-Hz ASSR. Inter-trial phase coherence maps of each sensor show the trial-to-trial phase-locking at 40 Hz.

## 4. Discussion

This was the first study to measure ASSR using OPM sensors to the best of our knowledge. We recorded the auditory brain activities using six OPM sensors inside of a two-layer magnetically shielded room. We observed the AEFs related to the pure-tone burst stimuli and the field distributions. We found obvious power changes and phase synchronization of ASSR modulated at 40 Hz.

In this study, we measured the AEFs using OPM sensors and confirmed that our results replicate the findings of previous OPM studies (Borna et al., 2020; Borna et al., 2017; Kim et al., 2014; Kowalczyk et al., 2021). We found clear M50 and M100 components of AEFs and obtained the topological map pattern. We further found that the M50 and M100 components had topologically opposite polarity. The brain source of the M50 component is known to be oriented toward the anterior/dorsal of the head whereas that of the M100 component is oriented toward the posterior/ventral of the head. These components were oriented approximately in opposite directions and had opposite topological patterns.

The latency and amplitude of the AEFs have been thought to be related to cognition and child development. The latencies of AEF components have been reported to be negatively related to age, reflecting neural development (An et al., 2020; Ponton et al., 2002; Ponton et al., 2000). The amplitudes of AEF components have been reported to be correlated to neural development and cognitive functions (An et al., 2020). In addition, the late latencies and weak amplitude of the AEF component have been shown in individuals with neurodevelopmental disorders, such as autism spectrum disorders (Roberts et al., 2010; Williams et al., 2021; Yoshimura et al., 2021). The investigation of AEFs is a potentially useful method of studying child development and neurophysiological disorders. To this end, the OPM sensor can be used in a flexibly fitting sensor array for small heads with only a small number of sensors over the region of interest of the brain area, and therefore, it is potentially an effective tool for child development and clinical research. The OPM sensor is compact and has flexible placement. These features allow personalized sensor arrangements according to the head size and head shape, as seen for EEG sensors. The OPM sensor can be used to measure the neuromagnetic field in a personalized position according to the head size and shape with a small number of sensors. Its use would therefore minimize the burden on the participant during recording, especially for child participants.

Our study showed the obvious power enhancement and phase synchronization of ASSR at 40 Hz. These results are consistent with the findings of previous EEG and MEG studies (Hari et al., 1989; Pastor et al., 2002; Tan et al., 2015). The auditory cortex is known to have a resonance frequency of approximately 40 Hz in humans (Galambos et al., 1981; Pastor et al., 2002; Ross et al., 2000). The 40-Hz ASSR is an evoked neural rhythm that is entrained by the external repeated auditory stimuli (Hari et al., 1989). The 40-Hz ASSR is thought to reflect the neural function of gamma-band-activities and neural inhibitory–excitatory balance (Sivarao et al., 2016; Tada et al., 2020). For this reason, previous studies have found that 40-Hz ASSR reduces power and phase synchronization in patients with schizophrenia (Tada et al., 2020; Thune et al., 2016), bipolar disorder (O’Donnell et al., 2013; Rass et al., 2010), and autism spectrum disorders (Ono et al., 2020; Seymour et al., 2020). The 40-Hz ASSR has high test–retest reliability (McFadden et al., 2014; Tan et al., 2015) and is thus thought to be useful in potential cognitive and clinical research. The neural synchronization of ASSR has been shown to undergo age-related changes in neural development (Cho et al., 2015; Edgar et al., 2016). The synchronization is therefore thought to be a useful indicator for research on child development and developmental disorders. The atomic signal gain of an OPM intrinsically depends on its detection frequency (Lee et al., 2014). Thus, the phase analysis of measured brain signals using an OPM is supposed to be questionable. However, as in the case of 40-Hz ASSR described in this article, a narrow-band analysis provides a reasonable phase synchronization result and power change. This study provides thus the groundwork for further neuronal phase analysis in OPM-MEG studies.

In this study, we obtained clear brain activities with a high signal-to-noise ratio even though we measured the OPM-MEG signals in a two-layer magnetically shielded room. This good signal-to-noise ratio might result from the close separation of the OPM sensors and source of the brain signals. OPM sensors can be placed near the scalp with a gap of approximately 3 mm because the OPM sensor operates at room temperature. The strength of the magnetic field is inversely proportional to the distance squared. OPM sensors therefore record clear signals with a high signal-to-noise ratio owing to their proximity to the signal source.

We recorded auditory brain activities using relatively few OPM sensors. We placed six OPM sensors so as to individually cover the T3 point with center sensor; the T3 point is the scalp site of EEG overlying the left-hemisphere auditory area of the cerebral cortex. We found that the auditory brain signals were strongest for the sensor over the T3 point. Steady-state responses recorded through EEG have been used for brain–computer interfaces (Ahn et al., 2015; Hill and Scholkopf, 2012; Muller-Putz and Pfurtscheller, 2008). Our OPM-MEG results open the door to developing practical brain–computer interfaces, which have thus far mainly relied on EEG recordings.

In this study, we measured the AEFs and 40-Hz ASSR using OPM sensors for healthy participants. We hope that our results will provide the groundwork for future OPM-MEG studies on child development, clinical practice, and brain–computer interfaces.

## Abbreviation

ASSR: auditory steady-state response
ERSP: event-related spectral perturbation
ITPC: inter-trial phase coherence
EEG: electroencephalography
MEG: magnetoencephalography
SQUID: superconducting quantum interference device
OPM: optically pumped magnetometer
AEF: auditory evoked field
TFR: time-frequency representation

## CRediT authorship contribution statement

**Kyung-min An:** Conceptualization, Methodology, Investigation, Formal analysis, Visualization, Writing – Original Draft, Project administration, Funding acquisition. **Jung Hyun Shim:** Conceptualization, Methodology, Resources, Funding acquisition. **Hyukchan Kwon:** Methodology, Software. **Yong-Ho Lee:** Methodology, Resources. **Kwon-Kyu Yu:** Methodology, Resources. **Moonyoung Kwon:** Investigation, Formal analysis, Visualization, Writing – Original Draft. **Woo Young Chun:** Resources, Funding acquisition. **Tetsu Hirosawa:** Conceptualization, Project administration. **Chiaki Hasegawa:** Conceptualization, Project administration. **Sumie Iwasaki:** Resources. **Mitsuru Kikuchi:** Conceptualization, Supervision, Funding acquisition. **Kiwoong Kim:** Conceptualization, Methodology, Visualization, Supervision, Writing – Original Draft, Project administration.

## Declaration of Competing Interest

All authors declare that they have no potential competing conflicts of interests.

## Acknowledgments

This work was supported by a grant from the Center of Innovation Program from the Japan Science and Technology Agency, the Collaborative Research Program of the Collaborative Research Network for Asian Children with Developmental Disorders, a grant from Korea Research Institute of Standards and Science (GP2021-0010), and the BK21 FOUR funded by the Ministry of Education (MOE) of Korea and National Research Foundation (NRF) of Korea. The funder had no role in the study design, data collection and analysis, decision to publish, or preparation of the manuscript. The authors thank Min-Young Kim for comment on the application to the institutional review board. They also thank Seong-mHwang, Seong-Joo Lee, Sangwon Oh, Jin-Mok Kim, Hyun Joon Lee, and Bogyung Kim for discussions on the setup of the OPM-MEG system.

## Data and code availability statements

The data supporting the findings of this study are available on request from the corresponding author, K.A. The data are not publicly available as they contain information that could compromise the privacy of the research participants.

## Notes

### Competing Interest Statement

The authors have declared no competing interest.

